# Co-released Norepinephrine and Galanin Act on Different Timescales to Promote Stress-Induced Anxiety-Like Behavior

**DOI:** 10.1101/2020.12.05.413138

**Authors:** Rachel P. Tillage, Stephanie L. Foster, Daniel Lustberg, L. Cameron Liles, David Weinshenker

**Author notes:** Corresponding author: David Weinshenker, Ph.D., Department of Human Genetics, Emory University School of Medicine, Whitehead 301, 615 Michael St., Atlanta, GA 30322.

## Abstract

**Background:** Both the noradrenergic and galaninergic systems have been implicated in stress-related neuropsychiatric disorders, and these two neuromodulators are co-released from the stress-responsive locus coeruleus (LC); however, the individual contributions of LC-derived norepinephrine (NE) and galanin to behavioral stress responses are unclear. Here we aimed to disentangle the functional roles of co-released NE and galanin in stress-induced behavior.

**Methods:** We used foot shock, optogenetics, and behavioral pharmacology in wild-type (WT) mice and mice lacking either NE (*Dbh^-/-^*) or galanin (*Gal^cKO-Dbh^*) specifically in noradrenergic neurons to isolate the roles of these co-transmitters in regulating anxiety-like behavior in the elevated zero maze (EZM) either immediately or 24 h following stress.

**Results:** Foot shock and optogenetic LC stimulation produced immediate anxiety-like behavior in WT mice, and the effects of foot shock persisted for 24 h. NE-deficient mice were resistant to the anxiogenic effects of acute stress and optogenetic LC stimulation, while mice lacking noradrenergic-derived galanin displayed typical increases in anxiety-like behavior. However, when tested 24 h after foot shock, both *Dbh^-/-^* and *Gal^cKO-Dbh^* mice lacked normal expression of anxiety-like behavior. Pharmacological rescue of NE, but not galanin, in knockout mice during EZM testing was anxiogenic. In contrast, restoring galanin, but not NE, signaling during foot shock normalized stress-induced anxiety-like behavior 24 h later.

**Conclusions:** These results indicate that NE and noradrenergic-derived galanin play complementary, but distinguishable roles in behavioral responses to stress. NE is required for the expression of acute stress-induced anxiety, while noradrenergic-derived galanin mediates more persistent responses following a stressor.

## INTRODUCTION

Stress is an important adaptive mechanism that allows animals to respond appropriately to threatening stimuli in their environment; however, stress is also a primary risk factor in the development of many neuropsychiatric disorders in humans, including anxiety and depression. These disorders are exceedingly common, with both major depressive disorder and all anxiety disorders having a lifetime prevalence rate of approximately 30% (1-3). Understanding the underlying neurobiological mechanisms that regulate adaptive and maladaptive behavioral responses to stress and how these mechanisms relate to psychopathology is necessary for the discovery of new, effective treatments for people suffering from anxiety and depression.

Both the central noradrenergic and galaninergic systems have been implicated in stress-related neuropsychiatric disorders. These two neuromodulators also share a neuroanatomical relationship, as galanin is co-expressed with norepinephrine (NE) in a majority of neurons in the locus coeruleus (LC), the primary source of NE in the brain, across species, including mice, rats, monkeys, and humans (4-9). Most LC research has focused exclusively on NE, despite the fact that galanin has also been independently implicated in stress-induced behaviors in rodents and stress-related neuropsychiatric disorders in humans (10-13). Stress strongly activates LC neurons, which broadly innervate the rest of the brain and play an important role in regulating attention, arousal, and the fight-or-flight response (14). Because the LC is a major source of both NE and galanin to many stress-responsive brain regions, it is a powerful neuroanatomical substrate for studying the contribution of NE and galanin co-transmission to stress-induced behavior. Neuropeptides, like galanin, have distinct dynamics from small molecule neurotransmitters like NE due to different mechanisms controlling synthesis, release, and degradation (15). Neuropeptides are thought to be preferentially released when neuronal firing rates are high, and previous research has suggested that galanin transmission occurs under conditions that strongly activate noradrenergic neurons, such as stress, highlighting the importance of understanding the individual contributions of galanin and NE from the LC in stress-induced behavior (15-17).

Traditional pharmacological and genetic studies allow functional investigation of single or even combinations of neurotransmitters, but they often lack the resolution to manipulate each one independently in a single cell type. On the other hand, application of modern neuroscience tools, such as optogenetics and chemogenetics, can achieve cell type-specificity but modulate neurotransmission indiscriminately, obscuring the potentially unique effects of co-released molecules. To overcome these limitations, we used genetically modified mouse lines that lack either NE (dopamine b-hydroxylase knockout; *Dbh^-/-^* mice) or galanin (*Gal^cKO-Dbh^* mice) specifically in noradrenergic neurons. Importantly, we previously showed that *Gal^cKO-Dbh^* mice have normal NE levels, and we show here that *Dbh^-/-^* mice have normal galanin expression in the LC (18), allowing for the isolation of each neuromodulator without affecting the other. *Dbh^-/-^* mice have been extensively studied since their initial creation 25 years ago (19), and do not differ from controls in traditional tests for anxiety-like behavior at baseline (20-22). Likewise, *Gal^cKO-Dbh^* mice also show normal anxiogenic behavior at baseline in classic anxiety assays, although have increased active coping behavior in some non-canonical paradigms, suggesting a role for NE-derived galanin in mediating passive coping behaviors (18). Here we used these two mouse lines to dissect the functional roles of co-released NE and galanin in regulating acute and persistent stress-induced anxiety-like behavior.

## METHODS AND MATERIALS

### Animals

All procedures related to the use of animals were approved by the Institutional Animal Care and Use Committee of Emory University and were in accordance with the National Institutes of Health guidelines for the care and use of laboratory animals. All mice were group housed, unless noted otherwise, and maintained on a 12/12 h light-dark cycle with access to food and water *ad libitum*. All manipulations and behavioral tests occurred during the light cycle. Balanced numbers of adult male and female mice (3-9 months) were used for all experiments. No sex differences were seen and data for males and females were combined for all analyses.

Because DBH is required for NE synthesis, *Dbh^-/-^* mice lack NE completely throughout the body. These mice were generated and maintained on a mixed 129/SvEv and C57BL/6J background, as previously described (19, 23) (see supplement for details). *Dbh^+/-^* littermates were used as controls in experiments with *Dbh^-/-^* mice because they are indistinguishable from WT (*Dbh^+/+^*) mice in behavior and catecholamine levels (19, 22, 24-26). *Gal^cKO-Dbh^* mice were generated as described previously and maintained on a C57BL/6J background by crossing *Dbh^cre+^;Gal^flx/flx^* mice to *Dbh^cre-^;Gal^flx/flx^* mice (18). Because of the difference in background strains, *Dbh^-/-^* mice could not be directly compared to *Gal^cKO-Dbh^* mice, so all experiments were designed to compared *Dbh^-/-^* mice to their *Dbh^+/-^* littermates, and *Gal^cKO-Dbh^* mice to their *Dbh^cre-^;Gal^flx/flx^* littermates (referred to as WT).

### RNAscope

Mice were briefly anesthetized and rapidly decapitated. Brains were collected, frozen in OCT, and stored at −80°C until sectioning. The LC was sectioned at 16-μm and slides were stored at −80°C. RNAscope was performed using the RNAscope Fluorescent Multiplex Assay (v1) according to manufacturer’s instructions (Advanced Cell Diagnostics, Newark, CA). Tissue was pre-treated as instructed using the RNAscope Sample Preparation and Pretreatment Guide for Fresh Frozen Tissue. The RNAscope assay was then performed according to manufacturer’s instructions using probes for galanin and tyrosine hydroxylase (TH). See supplement for imaging and quantification details.

### Stress paradigm

The foot shock stress protocol was conducted as described previously (27). Mice were individually exposed to 20 min of foot shock consisting of 19 shocks randomly interspaced by 30, 60 or 90 s (0.5 ms shocks, 1 mA) in chambers (Coulbourn Instruments, Holliston, MA) with an electric grid shock floor. Control animals were placed in the chamber for 20 min but were not administered shocks. Corticosterone (CORT) measurement was conducted as described (27) (see supplement for details).

### Optogenetics and EZM testing

Stereotaxic viral and optic ferrule surgeries were performed as described previously (27) (see supplement for details). LC optogenetic stimulation was based on a published paradigm used in our lab previously (27, 28). Photostimulation was delivered to the LC for 30 min (5 Hz tonic stimulation, 10 ms light pulses, 473 nm) in 3 min on/off bins in the home cage immediately before testing in the EZM, which was conducted as previous described (18). The EZM was used because it is a validated test for anxiety-like behavior in rodents and is sensitive to LC manipulations (28, 29). Furthermore, both mutant mouse lines show normal behavior at baseline in this assay, allowing us to observe stress-induced behavioral changes (18, 20). Histology in mice used for optogenetics was performed as described (27) (see supplement for details).

### Drugs

The nonspecific galanin receptor agonist galnon (Bachem, Torrance, CA, USA), which has a half-life of approximately 60 min (30), was dissolved in 0.9% sterile saline with 1% DMSO. Mice were injected with vehicle or galnon (2 mg/kg, i.p.) 20 min prior to either stress or behavioral testing.

The synthetic NE precursor L-3,4-dihydroxyphenylserine (DOPS) (Lundbeck, Deerfield, IL), was dissolved in distilled water with 2% HCl, 2% NaOH, and 2 mg/kg vitamin C, as described (21, 23). *Dbh^-/-^* mice were injected with vehicle or a cocktail of DOPS (0.5 g/kg, s.c.) and benserazide (250 mg/kg, s.c.) (Sigma-Aldrich), then tested 5 h later when NE levels peak (23, 31). Benserazide is an aromatic acid decarboxylase inhibitor that cannot cross the bloodbrain barrier and was used to prevent the conversion of DOPS to NE in the periphery, thus restricting restoration of NE to the brain (32).

### Statistical analysis

Data were found to be normally distributed using the D’Agostino-Pearson test and analyzed by one-way or two-way ANOVA, with Dunnett’s or Sidak’s correction respectively for multiple comparisons to examine planned comparisons between stress, virus, or drug treatment, where appropriate. Significance was set at p<0.05, and two-tailed variants of tests were used. Data are presented as mean ± SEM. Calculations were performed, and figures created using Prism Version 8 (GraphPad Software, San Diego, CA).

## RESULTS

### Galanin mRNA levels in the LC are normal in mice lacking NE

We previously showed that *Gal^cKO-Dbh^* mice have normal NE levels (18), but to isolate the role of each neuromodulator, we needed to ensure that galanin levels were unchanged in the LC of *Dbh^-/-^* mice. Using RNAscope to visualize galanin and TH mRNA transcripts in the LC (**Fig. 1a**), we found that there was no significant difference between the average galanin signal of *Dbh^-/-^* mice and littermate controls (t_6_=0.5583, p=0.6007) (**Fig. 1b**).

**Figure 1.**
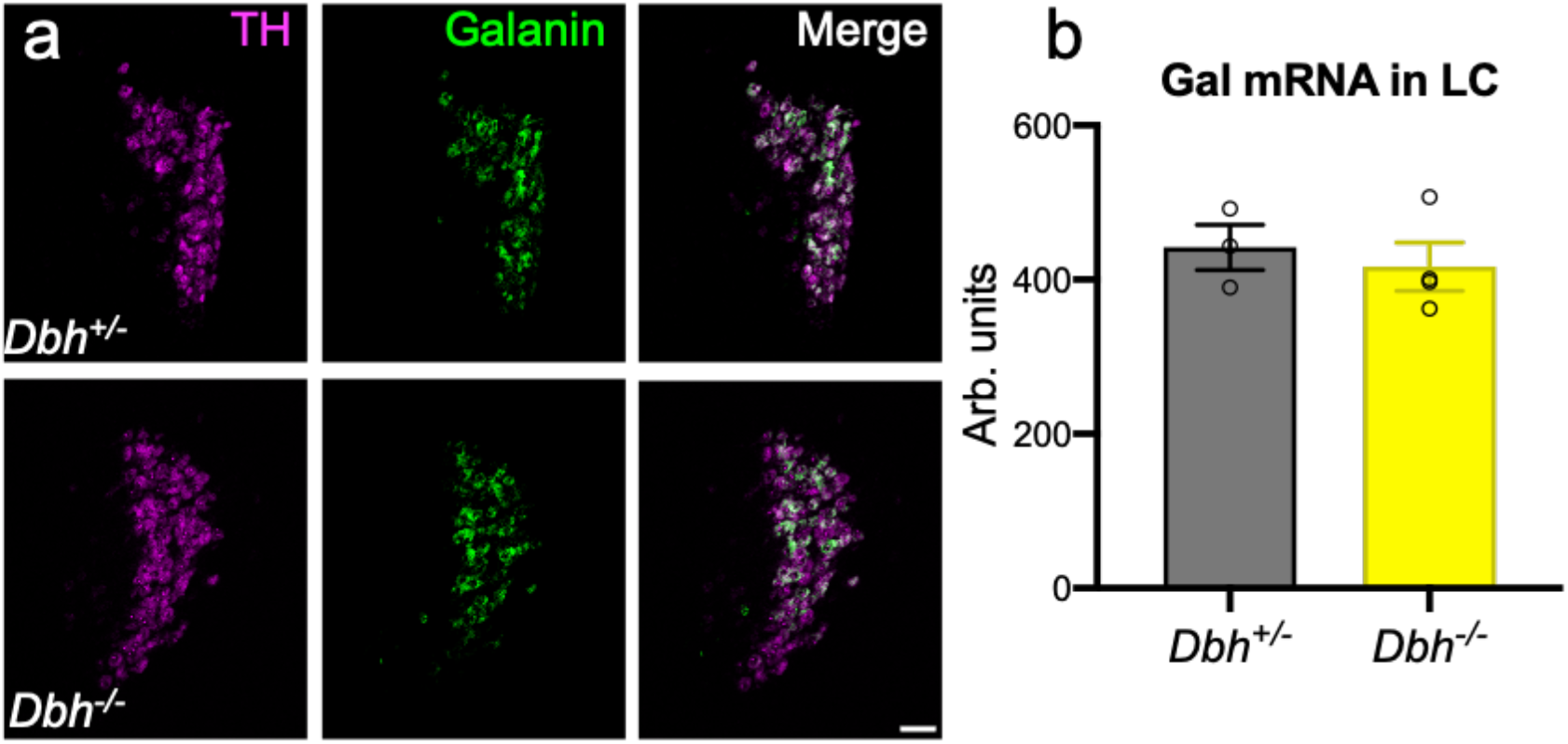
Galanin mRNA levels in the LC are normal in mice lacking NE. *Dbh^+/-^* control and *Dbh^-/-^* mice have equivalent levels of galanin mRNA in the LC as measured by average fluorescence intensity using RNAscope. (a) Representative images of TH (magenta) and galanin (green) mRNA expression and overlap in the LC from *Dbh^+/-^* and *Dbh^-/-^* mice (scale bar, 50 μm). (b) Quantification of galanin average fluorescence intensity. *n* = 4 mice per group, 2-4 LC sections per mouse. Error bars show SEM.

### Dbh^-/-^ mice, but not Gal^cKO-Dbh^ mice, are resistant to the acute anxiogenic effects of foot shock stress

To determine the relative importance of NE and noradrenergic-derived galanin for the acute behavioral response to a stressor, we subjected mice to 20 min of foot shock, and then immediately tested them in the EZM. When we analyzed the percentage of time the animals spent in the open segments of the EZM, a two-way ANOVA showed a significant genotype x treatment interaction for *Dbh^-/-^* mice compared to their *Dbh^+/-^* littermates (F_1,29_=8.003, p=0.0084). Post hoc tests revealed a significant decrease in percent time spent in the open segments for stressed *Dbh^+/-^* mice (t_29_=2.444, p=0.0413), but no difference between stressed and nonstressed *Dbh^-/-^* mice (t_29_=1.541, p=0.2503) (**Fig. 2a)**. For the *Gal^cKO-Dbh^* line, there was a main effect of treatment (F_1,30_=17.89, p=0.0002), but no genotype x treatment interaction (F_1,30_=0.00284, p=0.9580), indicating that foot shock stress decreased open segment exploration equally in WT and mutants (WT: t_30_=3.028, p=0.0100, *Gal^cKO-Dbh^*: t_30_=2.953, p=0.0121) (**Fig. 2b**).

**Figure 2.**
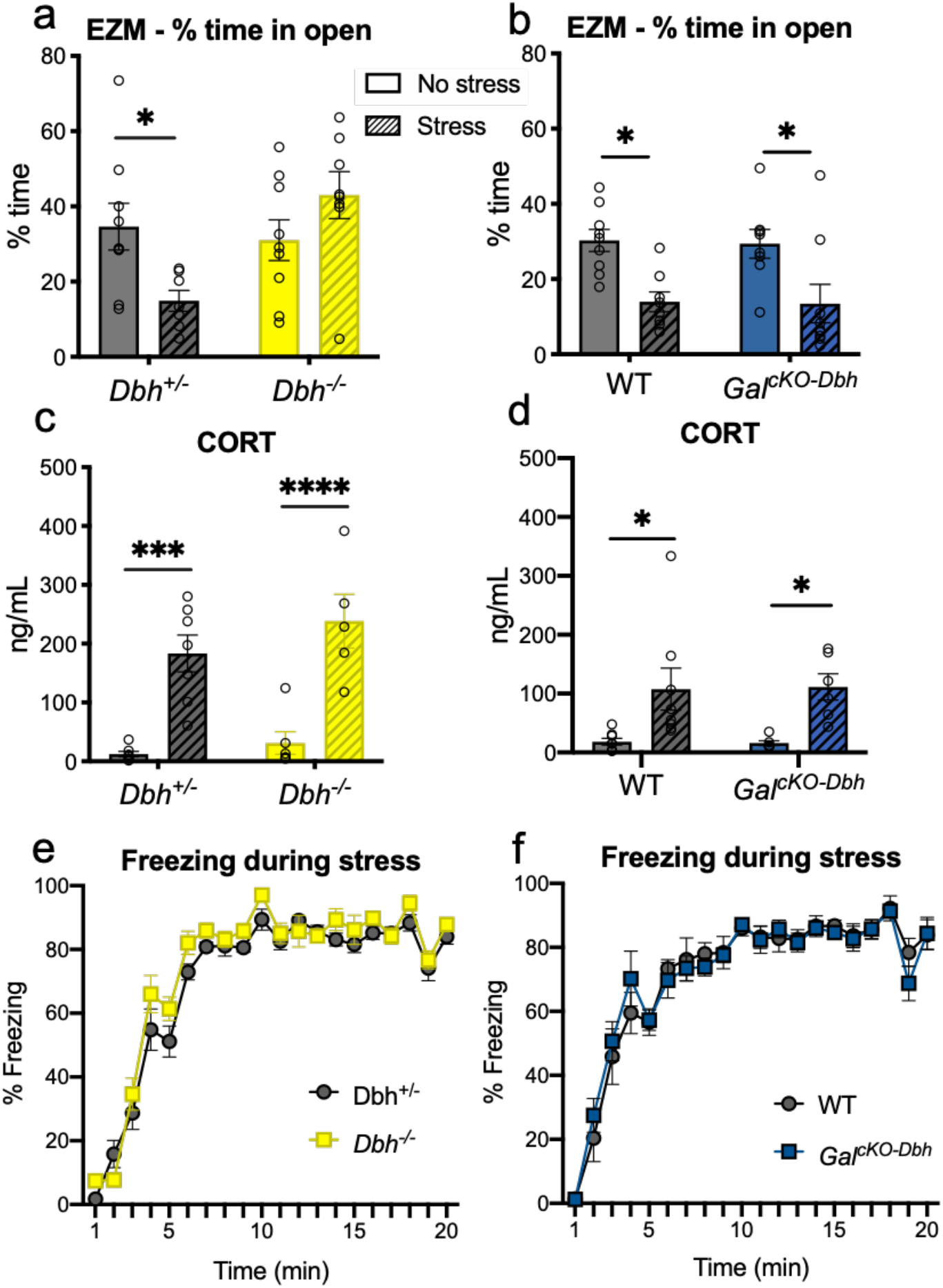
NE, but not noradrenergic-derived galanin, is required for anxiety-like behavior immediately following foot shock stress. *Dbh^-/-^, Gal^cKO-Dbh^*, and their respective littermate controls (*Dbh^+/-^* and WT) received 20 min of foot shock (“Stress”) or no foot shock (“No stress”) and were tested in the elevated zero maze (EZM) immediately afterwards. A separate group of mice received foot shock or no foot shock, and blood was collected immediately afterwards for CORT measurements. (a) *Dbh^+/-^*, but not *Dbh^-/-^* mice, showed a significant decrease in the percent time spent in the open segments of the EZM immediately after foot shock stress. (b) Both *Gal^cKO-Dbh^* and their WT littermates showed a significant decrease in the percent time spent in the open segments in the EZM immediately after foot shock stress. *Dbh^-/-^* and *Gal^cKO-Dbh^* mice showed increases in plasma CORT immediately following the foot shock stress (c-d) and similar freezing behavior during the foot shock stress (e-f) as their respective controls. *n* = 5-8 mice per group for CORT analysis. *n* = 8-9 mice per group for behavior. Error bars show SEM. *p<0.05, ***p<0.001, ****p<0.0001.

To confirm that the endocrine stress response was intact in mice lacking NE or NE-derived galanin, we measure levels of CORT in the blood immediately after exposure to foot shock stress. For the *Dbh^-/-^* mice, there was a significant effect of treatment on CORT levels (F_1,21_=48.49, p<0.0001), but no effect of genotype (F_1,21_=1.854, p=0.1877), indicating that both *Dbh^-/-^* mice and *Dbh^+/-^* controls have similar CORT responses to the foot shock stress (*Dbh^+/-^*: t_21_=4.759, p=0.0002, *Dbh^-/-^*: t_21_=5.088, p<0.0001) (**Fig. 2c**). Similarly, there was a significant effect of treatment on CORT levels for the *Gal^cKO-Dbh^* mice (F_1,24_=15.87, p=0.0005), but no effect of genotype (F_1,24_=0.001293, p=0.9716), indicating that *Gal^cKO-Dbh^* mice also have a normal CORT response to the foot shock stress (WT: t_24_=2.942, p=0.0142, *Gal^cKO-Dbh^*: t_24_=2.722, p=0.0237) (**Fig. 2d**). Both the *Gal^cKO-Dbh^* mice and the *Dbh^-/-^* mice also showed normal freezing to the foot shocks compared to their littermate controls during the stress itself (*Dbh^+/-^* vs. *Dbh^-/-^*: F_1,21_=3.690, p=0.0684; WT vs. *Gal^cKO-Dbh^:* F_1,10_=0.000936, p=0.9925) (**Fig. 2e-f**).

### Dbh^-/-^ mice, but not Gal^cKO-Dbh^ mice, show reduced acute anxiogenic effects of optogenetic LC stimulation

To complement the acute foot shock experiment and more selectively activate NE and galanin co-expressing neurons, we optogenetically stimulated LC neurons in WT, *Dbh^-/-^*, and *Gal^cKO-Dbh^* mice immediately before EZM testing (**Fig. 3a**). Mice received intra-LC infusion of virus expressing either ChR2-mCherry or mCherry alone driven by the noradrenergic-specific PRSx8 promoter, resulting in high levels of viral expression selectively in the LC. Viral expression and optic ferrule placement were confirmed in all animals by histology (**Fig. 3b**), and successful activation of the LC following optogenetic stimulation was confirmed via an increase in c-Fos immunoreactivity (*Dbh* mice: t_12_=5.055, p=0.0003; Gal mice: t_17_=3.277, p=0.0044) (**Fig. 3c-d**).

**Figure 3.**
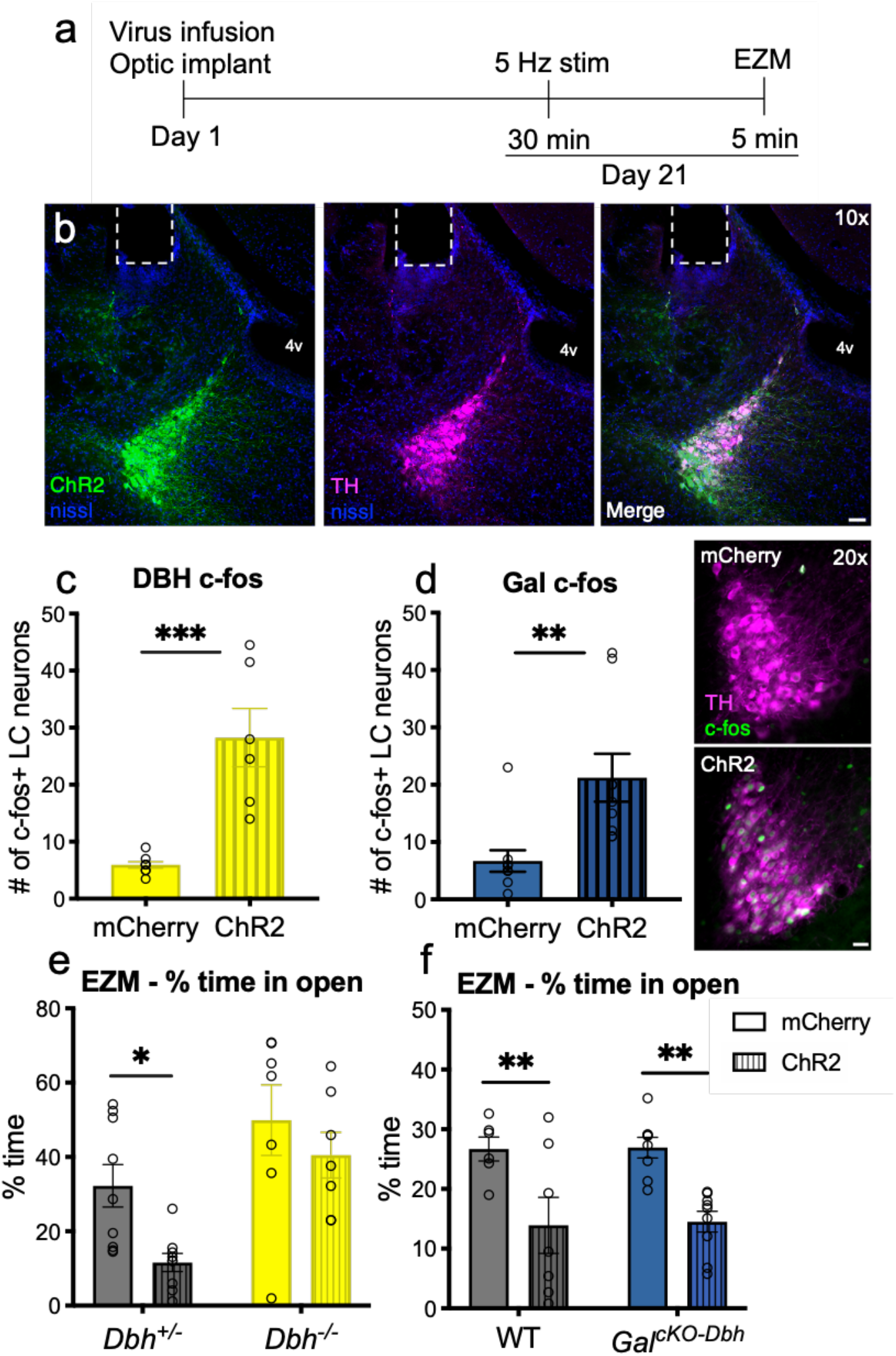
NE, but not noradrenergic-derived galanin, is required for anxiety-like behavior immediately following optogenetic LC stimulation. *Dbh^-/-^, Gal^cKO-Dbh^*, and their respective littermate controls (*Dbh^+/-^* and WT) expressing either ChR2-mCherry or mCherry alone in the LC were given 30 min of optogenetic LC stimulation (5 Hz tonic, 3 min on/off bins) and then tested in the elevated zero maze (EZM). At least one week following behavioral testing, mice were given 15 min optogenetic LC stimulation and brains were collected 90 min later for c-Fos immunohistochemistry. (a) Optogenetic surgery and behavior timeline. (b) Representative image of ChR2 expression (green) and tyrosine hydroxylase (TH, magenta) overlap in the LC and optic fiber ferrule placement (scale bar, 50 μm; 4v, 4^th^ ventricle). (c, d) Optogenetic stimulation of LC neurons expressing ChR2 resulted in greater c-Fos expression compared with LC neurons expressing mCherry. Because no differences in c-Fos expression between genotypes were observed, data were collapsed across genotype (TH, magenta; c-Fos, green; scale bar, 20 μm). (e) *Dbh^+/-^* mice expressing ChR2 spent less percent time in the open segments of the EZM following optogenetic simulation compared to mCherry-expressing *Dbh^+/-^* mice, while optogenetic LC activation did not impact the behavior of *Dbh^-/-^* mice. (f) Both *Gal^cKO-Dbh^* and their WT littermates expressing ChR2 showed a significant decrease in percent time spent in the open segments of the EZM following optogenetic stimulation compared to those expressing mCherry. *n* = 7-9 mice per group. Error bars show SEM. *p<0.05, **p<0.01.

Because no genotype differences in c-Fos expression after optogenetic stimulation were seen (data not shown), control and knockout animals were combined for c-Fos quantification for each line. For EZM behavior in the *Dbh^-/-^* mice, there were significant main effects of both virus (F_1,28_=6.108, p=0.0198) and genotype (F_1,28_=14.72, p=0.0006) on the percentage of time mice spent in the open segments, with planned comparisons showing that *Dbh^+/-^* ChR2 mice spent significantly less time in the open arms compared to *Dbh^+/-^* mCherry mice (t_28_=2.568, p=0.0314), whereas there was no significant difference between *Dbh^-/-^* mice expressing ChR2 or the control virus (t_28_=1.030, p=0.5263) (**Fig. 3e**). While a significant genotype x virus interaction was not seen in this experiment (F_1,28_=0.8574, p=0.3624), the lack of a significant decrease in time spent in the open segments of the EZM for the *Dbh^-/-^* ChR2 mice suggests that the NE is important for the full expression of optogenetic LC stimulation-induced anxiety-like behavior. For the *Gal* mice, there was a significant main effect of virus (F_1,26_=21.07, p<0.0001), but not genotype (F_1,26_=0.0238, p=0.8786), on the percentage of time spent in the open segments of the EZM (**Fig. 3f**), demonstrating that the noradrenergic galanin is dispensable for the normal behavioral response to optogenetic LC activation.

### Both Gal^cKO-Dbh^ and Dbh^-/-^ mice are resistant to the persistent anxiogenic effects of foot shock stress

Neuropeptides, such as galanin, have distinct dynamics compared to classic neurotransmitters like NE, and may therefore exert their effects on a different timescale. To examine the relative roles of NE and noradrenergic-derived galanin in the persistent behavioral response to a stressor, we subjected mice to 20 min of foot shock, and then tested them in the EZM 24 h later instead of immediately following the stressor (**Fig. 4a**). We found significant genotype x treatment interaction effects between stress and genotype in the percent time spent in the open segments for both *Dbh^-/-^* compared to *Dbh^+/-^* mice (F_1,38_=6.211, p=0.0172) (**Fig. 4b**) and *Gal^cKO-Dbh^* mice compared to WT controls (F_1,38_=4.694, p=0.0366) (**Fig. 4c**). Post hoc analyses revealed that the stress-exposed control animals in both experiments spent less time in the open segments of the EZM (*Dbh^+/-^*: t_38_=2.503, p=0.0332, WT: t_38_=2.556, p=0.0292), while *Dbh^-/-^* and *Gal^cKO-Dbh^* mice were resistant to this effect (*Dbh^-/-^*: t_38_=1.054, p=0.5081, *Gal^cKO-Dbh^*: t_38_=0.4633, p=0.8745). These results suggest that both neuromodulators are important for the long-term (24 h) anxiogenic-like effects of foot shock stress.

**Figure 4.**
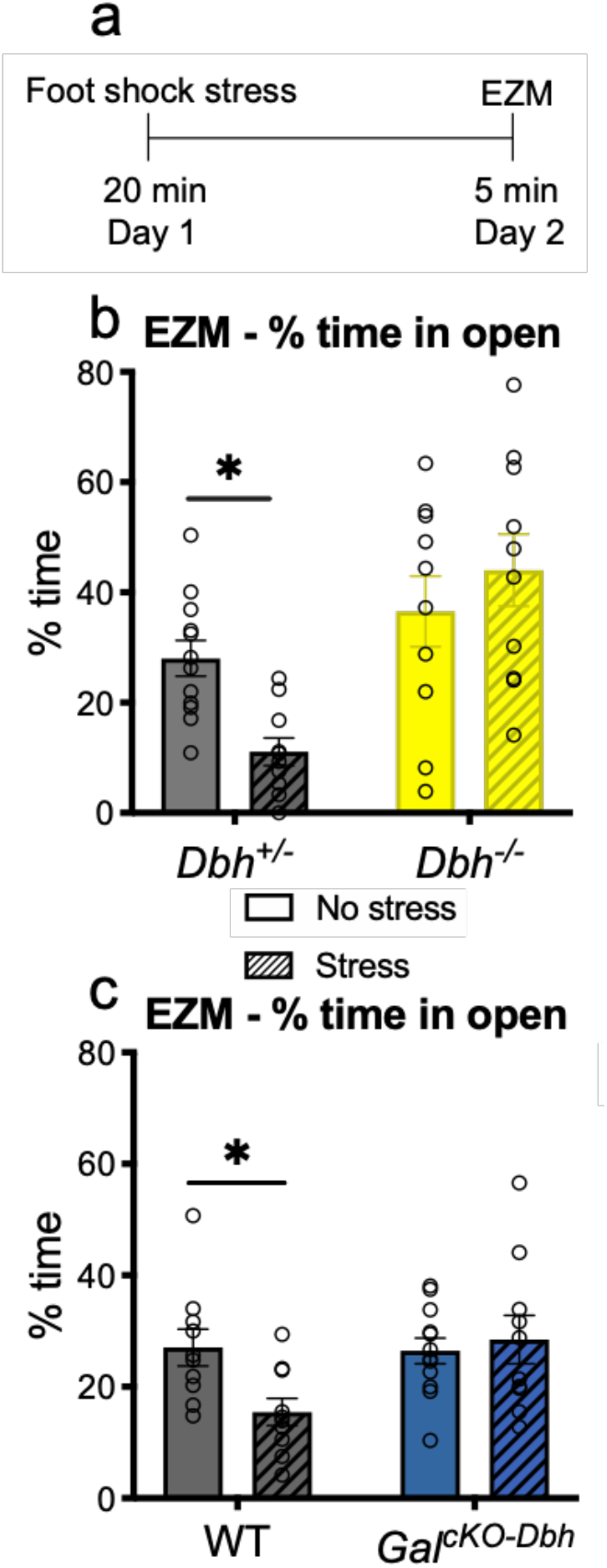
Both NE and noradrenergic-derived galanin are required for anxiety-like behavior 24 h following foot shock stress. *Dbh^-/-^, Gal^cKO-Dbh^*, and their respective littermate controls (*Dbh^+/-^* and WT) received 20 min of foot shock (“Stress”) or no foot shock (“No stress”) and were tested in the elevated zero maze (EZM) 24 h later. (a) Foot shock stress paradigm timeline. (b, c) *Dbh^+/-^* and WT, but not *Dbh^-/-^* or *Gal^cKO-Dbh^* mice, showed a significant decrease in percent time spent in the open segments of the EZM 24 h after foot shock stress. *n* = 10-12 mice per group. Error bars show SEM. *p<0.05, **p<0.01.

### Restoring galanin, but not NE, signaling during stress exposure rescues normal stress-induced anxiety-like behavior in knockout mice

To determine when galanin is acting to influence stress-induced anxiety-like behavior over 24 h, we injected *Gal^cKO-Dbh^* mice with the nonspecific galanin receptor agonist galnon to systemically increase galanin signaling either during the stressor or during the EZM 24 h later. We found no significant differences in percent time spent in the open segments of the EZM between stressed and non-stressed mice treated with galnon or vehicle during the EZM itself (F_1,27_=0.4267, p=0.8379) (**Fig. 5a**); however, when mice were treated with galnon prior to the foot shock, then tested in the EZM 24 h later, we found a significant interaction effect between stress and drug treatment (F_1,26_=4.238, p=0.0497), with post hoc analysis revealing a significant decrease in percent time spent in the open segments of the EZM for stressed *Gal^cKO-Dbh^* mice treated with galnon (t_26_=2.558, p=0.0331) (**Fig. 5b**). These results demonstrate that galanin transmission during the stressor leads to changes that impact stress-induced behavior 24 h later. In contrast, galanin signaling at the time of the behavioral test has no effect on EZM performance.

**Figure 5.**
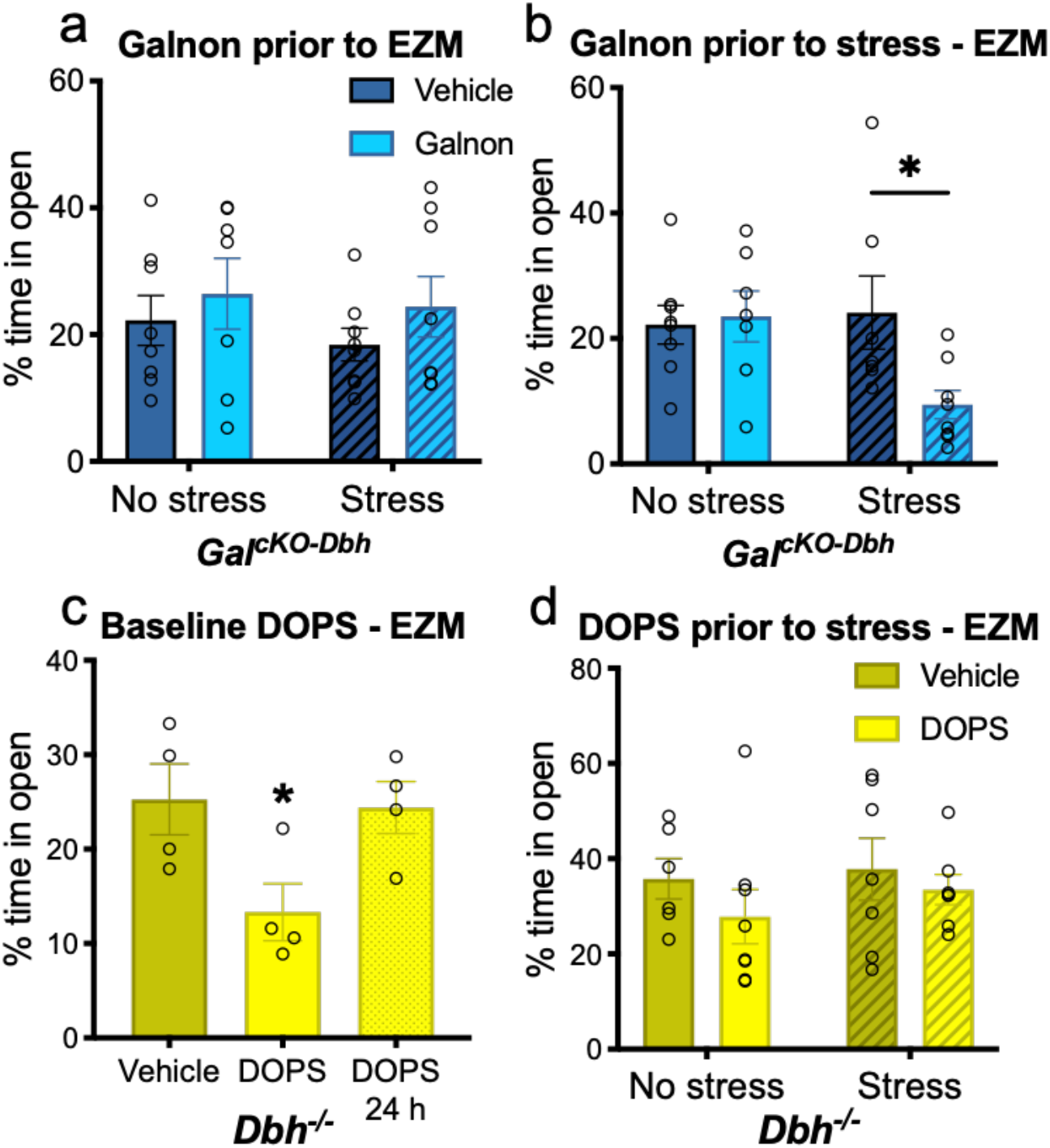
Restoration of galanin, but not NE, signaling during stress exposure rescues normal stress-induced anxiety-like behavior 24 h later. *Gal^cKO-Dbh^* and *Dbh^-/-^* were treated with the non-selective galanin receptor agonist galnon or the synthetic NE precursor DOPS, respectively. (a) Administration of galnon in *Gal^cKO-Dbh^* mice prior to EZM testing did not affect 24 h foot shock stress-induced anxiety-like behavior. (b) Galnon administration prior to the foot shock restored normal stress-induced anxiety-like effect 24 h later. (c) Acute DOPS treatment in *Dbh^-/-^* mice during EZM testing was anxiogenic compared to *Dbh^-/-^* mice treated with vehicle, but mice tested 24 h following DOPS administration displayed normal behavior. (d) Treatment of *Dbh^-/-^* mice with DOPS prior to foot shock stress exposure did not alter anxiety-like behavior 24 h later. *n* = 7-8 mice per group for a, b, d. *n* = 4 mice per group for c. Error bars show SEM. *p<0.05.

Previous research indicated that *Dbh^-/-^* mice display increased anxiety-like behavior following administration of the synthetic NE precursor DOPS in the elevated plus maze (21). Likewise, we found that pharmacological rescue of NE using DOPS in *Dbh^-/-^* mice during EZM testing was acutely anxiogenic compared to *Dbh^-/-^* mice treated with vehicle (F_2,9_=4.347, p=0.0477), while mice tested 24 h following DOPS had normal behavior (**Fig. 5c**). This indicates that the anxiogenic effects of acute NE restoration in *Dbh^-/-^* mice are transient and do not impact anxiety-like behavior 24 h later.

To determine whether NE in combination with stress can influence stress-induced anxiety-like behavior over 24 h, as we found with galanin, we administered DOPS to *Dbh^-/-^* mice to restore central NE signaling during foot shock stress, then tested the animals in the EZM 24 h later. There was no difference in percent time spent in the open segments of the EZM between stressed and non-stressed mice that had been treated with DOPS or vehicle during the stressor (F_1,24_=0.1201, p=0.7319) (**Fig. 5d**), suggesting that, unlike galanin, NE signaling during the stressor does not restore normal stress-induced anxiety-like behavior 24 h later in *Dbh^-/-^* mice.

## DISCUSSION

In this study, we aimed to resolve the roles of the LC co-transmitters NE and galanin in stress-induced behavior by using mice specifically lacking either NE or noradrenergic-derived galanin. This research is important because both of these transmitters regulate behavioral responses to stress in rodents and are implicated in stress-related neuropsychiatric disorders in humans (10-13). While both *Dbh^-/-^* and *Gal^cKO-Dbh^* mice responded similarly during the stressor itself, with high levels of freezing and increased plasma CORT, we found that that *Dbh^-/-^* mice were resistant to both foot shock and optogenetic LC activation-induced anxiety-like behavior in the EZM immediately after the manipulation, whereas the responses of *Gal^cKO-Dbh^* mice were similar to controls; however, when tested 24 h after foot shock stress exposure, we found that both *Dbh^-/-^* and *Gal^cKO-Dbh^* mice were resistant to the persistent effects of foot shock in EZM performance. Furthermore, we found that pharmacological restoration of galanin, but not NE, in knockout mice during foot shock stress normalized expression of stress-induced anxiety-like behavior 24 h following stress. In contrast, rescue of NE signaling in *Dbh^-/-^* mice alone at the time of EZM testing induced anxiety-like behavior. These results suggest that NE transmission is important for anxiogenic responses at the time of behavioral testing, but is dispensable during the stressor itself, while NE-derived galanin plays no acute role but rather is released during the stressor to promote anxiety-like behavior over the course of hours to days.

Our findings that the *Dbh^-/-^* mice were resistant to stress-induced behavioral changes add to growing literature supporting an important role for NE in mediating anxiogenic responses. The inability of NE restoration during foot shock to rescue normal anxiety-like behavior 24 h later in the *Dbh^-/-^* mice suggests that acute NE signaling at the time of the behavioral test is required. In support of this idea, administration of DOPS to *Dbh^-/-^* alone, in the absence of foot shock, reduced open segment exploration in the EZM (this study) and the related elevated plus maze (21). We speculate that because *Dbh^-/-^* mice lack NE from birth and have a compensatory upregulation of b-adrenergic receptors (βAR) (33), acute NE restoration has anxiogenic properties in the knockouts akin to cocaine-induced increases in WT animals (21). Recent studies have found that LC projections to the basolateral amygdala (BLA) mediate acute foot shock stress-induced increases in BLA firing through βAR signaling, and local administration of βAR antagonists in the BLA immediately following foot shock leads to decreased freezing behavior in an immediate fear extinction paradigm (34, 35). Furthermore, LC-NE-mediated βAR transmission in the BLA induced anxiety-like behavior on its own and modulated pain-related anxiety-like behavior (36, 37). This acute action of NE in the BLA likely explains the acute stress-induced anxiety-like effect in the EZM immediately following foot shock that we observed in WT and *Gal^cKO-Dbh^* mice, which is absent in the *Dbh^-/-^* mice.

A previous study showed that selective optogenetic activation of galanin-containing LC neurons was sufficient to induce avoidance behavior and this behavioral effect could be prevented by a bAR antagonist, indicating a noradrenergic mechanism (28); however, the potential contribution from galanin was not explicitly examined. Here we employed the same optogenetic protocol to selectively activate LC neurons in *Gal^cKO-Dbh^* mutants and observed increased anxiety-like behavior in the EZM similar to WT animals, but this effect was absent in *Dbh^-/-^* mice, which had no change in EZM performance. Our results support the previous work by McCall and colleagues by showing that acute optogenetic LC stimulation-induced anxiety-like behavior requires NE and expand on this finding by showing that NE-derived galanin is dispensable for this acute behavioral change.

Neuropeptides, such as galanin, have different dynamics than small molecule neurotransmitters like NE due to distinct mechanisms controlling synthesis, release, and degradation of the molecules (15). Because of their storage in large, dense core vesicles (LDCV), neuropeptides are thought to be preferentially released when neuronal firing rate and Ca^2+^ influx are high, and previous research has suggested that galanin transmission occurs under conditions that strongly activate noradrenergic neurons (15-17). Once released, galanin can act as an inhibitory neuromodulator by signaling acutely through GalR1 and GalR3, which are typically Gi-coupled, but galanin can also act as a neurotrophic factor via GalR2-Gq signaling, leading to changes in neuronal plasticity over time (12). Our results show that restoration of galanin signaling during the stressor, but not 24 h later during the EZM, can normalize stress-induced anxiety-like behavior in the *Gal^cKO-Dbh^*, strongly suggest that galanin is acting through a neurotrophic mechanism in this paradigm to establish persistent anxiety-like behavior. This mechanism could involve changes in dendritic spines or receptor distributions in downstream, stress-sensitive brain regions (e.g. amygdala, prefrontal cortex (PFC)). Alterations in plasticity take time to develop and thus would not change behavior in the EZM test until hours or days after the stress, and would fail to occur in the absence of galanin transmission from the LC in the *Gal^cKO-Dbh^* mice. Previous research has shown that significant atrophy and remodeling of dendrites occurs in stress-sensitive regions, such as the PFC, as early as 24 h after a single foot shock stress, and that chronic intracerebroventricular infusions of galanin prevent foot shock-induced dendritic spine loss in the PFC (38-40). Future experiments will be required to identify the mechanism underlying the persistent effects of stress-induced galanin transmission, including determining where galanin is acting in the brain and which receptor mediates its effects.

There are limitations associated with our study. While there are significant benefits to using knockout models like the *Dbh^-/-^* and *Gal^cKO-Dbh^* mice, it is important to note that although these genetic manipulations are specific to noradrenergic neurons, neither is specific to the LC. The LC certainly projects widely to brain regions known to control behavioral responses to stress (e.g., hippocampus, amygdala, PFC), and most LC neurons co-express galanin, making it the most likely source of galanin and NE that mediates the anxiogenic effects of stress; however, A2 noradrenergic neurons in the nucleus of the solitary tract also mediate anxiety-like behavior, although NE transmission from these cells appears to be anxiolytic rather than anxiogenic (41), and they express little if any galanin (18, 42). Because the knockout mice used here lack expression of NE or galanin throughout development, it is possible that compensatory mechanisms alter the way these systems function, although our findings that acute pharmacological restoration of NE and galanin signaling rescues stress responses in the *Dbh^-/-^* and *Gal^cKO-Dbh^* mice, respectively, seem to rule out that caveat in the present experiments. Finally, our finding that NE-derived galanin is required for the persistent anxiogenic effects of stress may appear at first blush to directly contradict our recent report that overexpression of galanin in noradrenergic neurons has the opposite effect and promotes stress resilience (27). Nonetheless, this inconsistency in the valence of how NE-derived galanin modulates stress resilience is consistent with the complexity often reported in the galanin literature, and potentially aligns with contrasting roles that galanin plays across different mechanisms (such as engagement of different galanin receptors in different brain regions) and timescales (hours to days versus weeks to months) (12).

In conclusion, our findings demonstrate that both NE and NE-derived galanin are involved in mediating aspects of stress-induced anxiety-like behavior and appear to act over different timescales. More broadly, our results may also shed light on how other small molecule and peptide co-transmitter systems function throughout the nervous system. Although some of the complexity created by co-transmission has been dissected with the aid of modern neuroscience techniques, such as using RNAi to knockdown individual co-transmitters in a single cell type (43), this remains an understudied area of research and there is much to discover about the functional effects of co-released neurotransmitters. Our findings on the differential roles of NE and galanin in the LC-NE system point to the importance of examining the roles of co-transmitters over different timescales and may be applicable to other neuromodulatory systems that co-express neuropeptides, such as the serotonergic dorsal raphe neurons which express neuropeptide Y and galanin in some species, or the dopaminergic ventral tegmental area neurons which express high levels of brain-derived neurotrophic factor (44, 45). Future studies will continue to disentangle the functional relationship between these co-expressed neuromodulators in the LC-NE system and determine the mechanism by which galanin regulates persistent adaptive and maladaptive stress responses, with the ultimate goal of leveraging this knowledge to improve treatments for stress-related disorders.

## ACKNOWLEDGEMENTS AND DISCLOSURES

This work was supported by the National Institutes of Health (NIH) (F31MH116622 to RPT, F31DA044726 to SLF, T32NS007480 to DL, R01NS102306, R01DA038453, R01DA049257, and R01AG061175 to DW). RPT and DW conceived, designed, and supervised the project. RNAscope *in situ* hybridization was performed by SF. Immunohistochemistry experiments were performed by RPT and DL. All behavioral experiments were performed and analyzed by RPT. Pharmacological experiments were performed by RPT with assistance from DW. Mouse husbandry and genotyping were performed by LCL. RPT and DW wrote the manuscript with input from co-authors. We thank C. Strauss for helpful editing of the manuscript. The authors report no financial interests or potential conflicts of interest.

