## Supplemental Methods for "Co-released Norepinephrine and Galanin Act on Different Timescales to Promote Stress-Induced Anxiety-Like Behavior"

**SUPPLEMENT**

**Materials and Methods**

### *Dbh^-/-^ mice breeding*

### To prevent embryonic lethality associated with the homozygous *Dbh* deficiency, pregnant *Dbh^+/-^* dams were given the adrenergic receptor agonists isoproterenol and phenylephrine (20 µg/ml each) from E9.5 to E14.5, and the synthetic precursor to NE, L-3,4-dihydroxyphenylserine (DOPS; 2 mg/ml), from E14.5 to parturition in their drinking water. Thus, *Dbh^-/-^* mice lack NE from birth.

### *RNAscope Imaging and quantification*

### After the final wash, slides were counterstained with DAPI and coverslipped with Prolong Diamond Antifade Mountant (Thermo Fisher Scientific, Waltham, MA). All RNAscope slides were imaged within 24-48 hours of performing the assay. Slides were imaged using a Nikon A1R HD25 confocal microscope with NIS Elements Software. For each slide, the LC was centered in the field of view and a 20x objective lens was used to acquire a 1024 x 1024 pixel single plane image taken at the focal plane with maximal probe intensity. Gain settings were pre-validated with positive and negative control probe slides and saved as an optical configuration that was used for all subsequent imaging. For quantification of galanin mRNA fluorescence intensity, 2-4 LC sections from each mouse were analyzed and averaged together to determine the mean for each subject. All image analysis was performed with ImageJ software for threshold application (Otsu method) and quantification. TH immunoreactivity was used to define LC area and only signal within that area was used for galanin mRNA signal quantification. Experimenter was blind to genotype during image collection and analysis.

### *Corticosterone (CORT) measurement*

### Mice used for CORT measurement went through the foot shock stress paradigm described in the regular Methods section, and were anesthetized with isoflurane 15 min later. Mice were rapidly decapitated and trunk blood was collected in EDTA-coated tubes (Sarstedt Inc., Newton, NC) and chilled on ice. Blood was centrifuged for 20 min at 3000 rpm at 4°C, and resulting plasma was collected and stored at -80°C. CORT was measured using the Enzo Life Sciences kit (Farmingdale, NY) following manufacturer’s small volume protocol for blood plasma, including diluting samples 1:40 with a steroid displacement reagent solution.

### *Stereotaxic surgery*

### For optogenetic experiments, mice were anesthetized with isoflurane and given the analgesic meloxicam (5 mg/kg, s.c.) at the start of the surgery. A lentiviral vector containing the channelrhodopsin (ChR2) construct with an mCherry tag under control of the noradrenergic-specific PRSx8 promoter (25) was infused unilaterally into the LC (-5.4AP; +1.2ML; -4.0DV) with a 5 µL Hamilton syringe and Stoelting Quintessential Stereotaxic Injector pump at a rate of 0.15 µL/min. Unilateral LC stimulation is sufficient to produce behavioral effects in mice that are indistinguishable from bilateral stimulation (26, 27). Control mice received a lentivirus containing mCherry alone under the PRSx8 promoter. Each animal received a 0.7 µL infusion, and the infusion needle was left in place for 5 min after infusion to allow for viral diffusion. An optic fiber ferrule (ThorLabs, Newton, NJ) was implanted 0.5 mm dorsal to the viral injection site (-5.4AP; +1.2ML; -3.5DV), and permanently attached to the skull with screws and dental acrylic. Mice were singly housed after this surgery to prevent cage mates from damaging the headcap and given at least 3 weeks to recover and allow for full viral expression before testing. Mice were habituated to handling and connection of the optic patch cable to the implanted optic ferrule for one week prior to testing.

### *Histology*

### Mice used for optogenetic experiments were exposed to 5 Hz LC photostimulation for 15 min in the home cage, then anesthetized 90 min later with isoflurane and transcardially perfused with potassium phosphate-buffered saline (KPBS), followed by 4% PFA in PBS. Brains were postfixed overnight by immersion in 4% PFA at 4°C, and then transferred to 30% sucrose in KPBS for 48 h at 4°C. Brains were flash frozen and tissue was cryosectioned at 40-µm. Viral expression and correct optic fiber targeting were assessed by immunostaining for the mCherry tag using rabbit anti-DsRed primary antibody (1:1000, #632496; Takara Bio) and Alexa Fluor 568 goat anti-rabbit secondary (1:500, A-11011; Invitrogen). Sections were co-stained for the noradrenergic marker tyrosine hydroxylase (TH) with chicken anti-TH (1:1000, AB76442; Abcam) and Alexa Fluor 488 goat anti-chicken secondary (1:500, AB150169; Abcam). In adjacent LC sections, activated LC neurons were detected with rabbit anti c-Fos primary antibody (1:5,000, ABE457; Millipore Sigma) with Alexa Fluor 488 goat anti-rabbit secondary (1:500, A-11008; Invitrogen), and TH-expressing cells were co-stained using chicken anti-TH (1:1000, AB76442; Abcam) with Alexa Fluor 568 goat anti-chicken secondary (1:500, A-11041; Invitrogen). After staining, all sections were mounted on slides and cover slipped with Fluoromount plus DAPI (Southern Biotech). Images were collected on a Leica DM6000B epifluorescent upright microscope at 10x or 20x. Mice were excluded from analyses if they did not show viral expression in the LC, correct optic fiber targeting, and increased c-Fos expression in the LC compared to the control animals of the same cohort. No differences in c-fos expression between genotypes were observed.
